# Single-orientation Quantitative Susceptibility Mapping identifies the Ventral Intermediate Nucleus of the Thalamus

**DOI:** 10.1101/2021.07.02.450973

**Authors:** Fang Frank Yu, Dongyoung Lee, Michael Achilleos, Fabricio Feltrin, Bhavya R. Shah

**Affiliations:** Neuroradiology and Neuro-intervention Section, Department of Radiology, University of Texas Southwestern Medical Center, Dallas, TX, United States; Advanced Imaging Research Center, University of Texas Southwestern Medical Center, Dallas, TX, United States; O’Donnell Brain Institute, University of Texas Southwestern Medical Center, Dallas, TX, United States; Department of Radiology, University of Texas Southwestern Medical Center, Dallas, TX, United States; Transcranial Focused Ultrasound Lab, University of Texas Southwestern Medical Center, Dallas, TX, United States; Department of Neurological Surgery, University of Texas Southwestern Medical Center, Dallas, TX, United States; Center for Alzheimer’s and Neurodegenerative Disease, University of Texas Southwestern Medical Center, Dallas, TX, United States

**Keywords:** MRI, MRI-guided therapy, Tremor, quantitative susceptibility mapping

## Abstract

**Introduction:** The ventral intermediate nucleus (VIM) represents the primary target in the treatment of tremor. Accurate localization is extremely important given its proximity to other thalamic nuclei. We utilized single orientation quantitative susceptibility mapping (QSM) at 3T to directly visualize the VIM.

**Methods:** Four adult volunteers, one adult cadaver, and an essential tremor patient were scanned on a 3T MRI using a multi-echo gradient echo sequence. QSM images were generated using the improved sparse linear equation and least-squares (iLSQR) algorithm. Two adult subjects underwent multiple head orientation imaging for multi-orientation QSM reconstruction. The VIM was prospectively identified with direct visualization as well as indirect landmark-based localization.

**Results:** The bilateral VIM was consistently identified as a hypointense structure within the lateral thalamus, appearing similar on multi-orientation and single-orientation QSM, corresponding to the myelinated dentatorubrothalamic tract (DRTT). The indirect method resulted in a comparatively inferomedial localization, at times missing the VIM and DRTT.

**Conclusion:** Single-orientation QSM offers a clinically feasible, non-invasive imaging-based approach to directly localize the VIM.

## Introduction

Tremor is the most common movement disorder, and it imposes a significant burden on quality of life. Tremor is most commonly seen in the setting of essential tremor and tremor predominant Parkinson’s disease (TDPD). First-line therapy for tremor is medical management but up to 30% of patients develop adverse effects from medications or the tremor becomes medically refractory. In these situations, the tremor can either be treated with surgical procedures such deep brain stimulation or radiofrequency thalamotomy. More recently, a non-invasive procedure, MR-guided high-intensity focused ultrasound (MRgHIFU), has been shown to effectively treat tremor.(1) MRgHIFU is currently FDA approved to treat ET and TDPD.

The target, the ventral intermediate (VIM) nucleus of the thalamus, is situated along the dentatorubrothalamic tract (DRTT), and is responsible for receiving and sending projections between the primary motor cortex (M1) and cerebellum. Loss of Purkinje cells in the cerebellum result in aberrant electrical impulses in the DRTT.(2) The resultant imbalance between inhibitory and excitatory pathways results in tremor. The VIM is not visible on standard, high-resolution MRI sequences. Therefore, indirect targeting based on anatomic landmarks is performed. Unfortunately, anatomic variability cannot be adequately accounted for with this approach. During Deep Brain Stimulation, micro-electrode recording can be used to identify neurons exhibiting tremor activity, but this increases cost and operating room times. Recently, advanced MRI techniques have been shown to identify the target (3, 4). Susceptibly sensitive imaging techniques, such as susceptibility weighted imaging (SWI), have shown the ability to delineate subtle contrast differences between brain structures.(5) The advent of quantitative susceptibility mapping (QSM) has overcome key limitations in using phase images, which is directly correlated to magnetic susceptibility of brain tissues.(6) Although both myelin and iron content contribute to the magnetic susceptibility signal, they do so in an opposing fashion. As a result, the two substrates are readily distinguishable.

Previous studies have shown that QSM can delineate deep grey nuclei such as the subthalamic nucleus.(7) Desitung et al. utilized multi-orientation QSM at 7T to delineate thalamic nuclei.(8) Historically, multi-orientation QSM is capable of producing higher quality images when compared to single-orientation methods.(9) However, scanning at multiple head orientations significantly increases scan time and contributes to patient discomfort. Furthermore, 7T MR scanners are not widely available. Our study seeks to overcome these limitations by using single-orientation QSM acquired at 3T to delineate the VIM. We further establish the generalizability of this approach across different MR scanner platforms with different scanning parameters.

## Methods

### Image Acquisition

This prospective study was approved by the IRB at the University of Texas Southwestern Medical Center. All seven living research participants underwent informed consent. MR imaging was performed on 3T MRI units (Siemens Prisma, Philips Ingenia, and Philips Achieva). Four healthy adult volunteers and one patient with essential tremor underwent imaging with a single head orientation multi-echo GRE protocol, and two healthy adult volunteers were imaged with multiple head orientation multi-echo GRE protocol. An overview of the different imaging parameters used in this study are shown in Supplementary Material Table S1.

### Image Processing

The phase data from the multiple echoes was unwrapped using Laplacian-based unwrapping and the normalized phase was subsequently calculated as previously described. The magnitude images were then fed through the Brain Extraction Tool in FSL (fsl.fmrib.ox.ac.uk) to generate brain masks. Background phase removal was then performed using the variable kernel sophisticated harmonic artifact reduction for phase data method (V-SHARP) with a spherical mean radius increasing from 0.6 mm at the boundary of the brain to 25 mm towards its center.(10) Single orientation QSM images were generated from the processed phase images using the improved sparse linear equation and least-squares (iLSQR) algorithm.(11) For the two subjects who underwent multi-orientation GRE imaging, COSMOS reconstruction was performed using an iterative least-squares formulation.(12)

### Localization of the VIM

The processed magnetic susceptibility maps were displayed within −0.05 to 0.15 ppm contrast window range and manually inspected by 2 board-certified neuroradiologists (F.Y. and B.R.S., 17 years of cumulative experience) to delineate the location of the VIM. Localization was based on visualization of a hypointense band within the mid thalamus. The VIM corresponds to the lateral 1/3^rd^ of the hypointense band, medial to the posterior limb of the ipsilateral internal capsule.

### Indirect Localization of the VIM

In order to streamline the calculation and visualization of the coordinates of the Guiot parallelogram, a customized script was developed in MATLAB 2019a (MathWorks, Natick, MA) with the SPM12 software toolkit (Wellcome Trust Centre for Neuroimaging, London, UK). After the subject’s 3D T1w brain MRI volume is loaded into the software using the SPM12 toolkit, the volume is displayed in three separate windows in orthogonal views (axial, coronal, sagittal) in standard radiographic conventions. The width of the third ventricle, the height of the thalamus, and the distance between the anterior commissure [AC] and posterior commissure [PC] are manually specified by the user, the locations of the four corners of Guillot’s parallelogram (C2, C3, C4, and C5) are determined in the real-world coordinate system (in mm), defined as follows. The reference system is centered on the center of the PC with the Y-axis passing through the AC with the positive Y-axis directed anteriorly. The (Y, PC, Z) plane is the median sagittal plane, with the positive Z pointing downward. The X-axis is perpendicular to the (Y, PC, Z) plane with the positive X-direction towards to the left side of the cadaver. The geometric definitions of the Guiot parallelogram are used to derive the equations for determining the coordinates per Dormont et al ^9^. Conversion of the coordinates within MRI volumes between voxel and real-world systems for displaying and calculation purposes, was achieved through the 4-by-4 affine transformation matrix read from each MRI volume via the SPM library.

### Atlas-based delineation of the thalamic nuclei

Anatomic structures, including the thalamic nuclei, were identified using a histoarchitectural atlas (http://nist.mni.mcgill.ca/?p=1209)(13, 14). The atlas dataset consisted of a population-averaged brain atlas with T1-weighting at 1 mm^3^ isotropic resolution, as well as a histology-derived digitized atlas containing 123 anatomical structures from the Schlatenbrand and Wahren atlas. Both the T1w brain atlas and the histology-derived atlas were in the same anatomic space (ICBM152). The T1w atlas images as well as the first echo from each subject’s GRE magnitude data were skull-stripped utilizing a modified Advanced Normalization Tools (ANTs) script.(15) Afterwards, the T1w atlas was co-registered to the GRE magnitude image using niftyreg (http://cmictig.cs.ucl.ac.uk/wiki/index.php/NiftyReg). The resultant transformation matrix was then applied to the histology-derived atlas for coregistration of the atlas to the QSM images for each individual subject.

### Comparison of QSM with Indirect Targeting and DRTT

The isocenter of the VIM determined based on direct visualization was compared to indirect targeting using the Guiot parallelogram, with the distance between the two sets of points calculated for both left and right VIM. For subject #6, tractography was performed using DTI data acquired after MR guided HIFU. DTI data was preprocessed with motion and distortion correction using Brainlab Elements Software (version 4.0). Deterministic tractography was then performed using fiber assignment by continuous tracking and tensor deflection methods. Isolation of the DRTT was performed as previously described (4). The patient’s processed QSM image was uploaded to Brainlab and co-registered to the subject’s DTI b_0_ image.

## Results

Several thalamic nuclei could be directly visualized on the quantitative susceptibility maps based on differences in magnetic susceptibility (Figure 1). Relatively iron rich nuclei such as the pulvinar and ventral anterior nucleus appeared as hyperintense structures. To produce adequate image contrast for visualization of the thalamic substructures, the images were windowed to between −0.07 and 0.15 ppm to allow sufficient contrast range to capture typically occurring diamagnetic and paramagnetic soft tissues.

**Figure 1:**
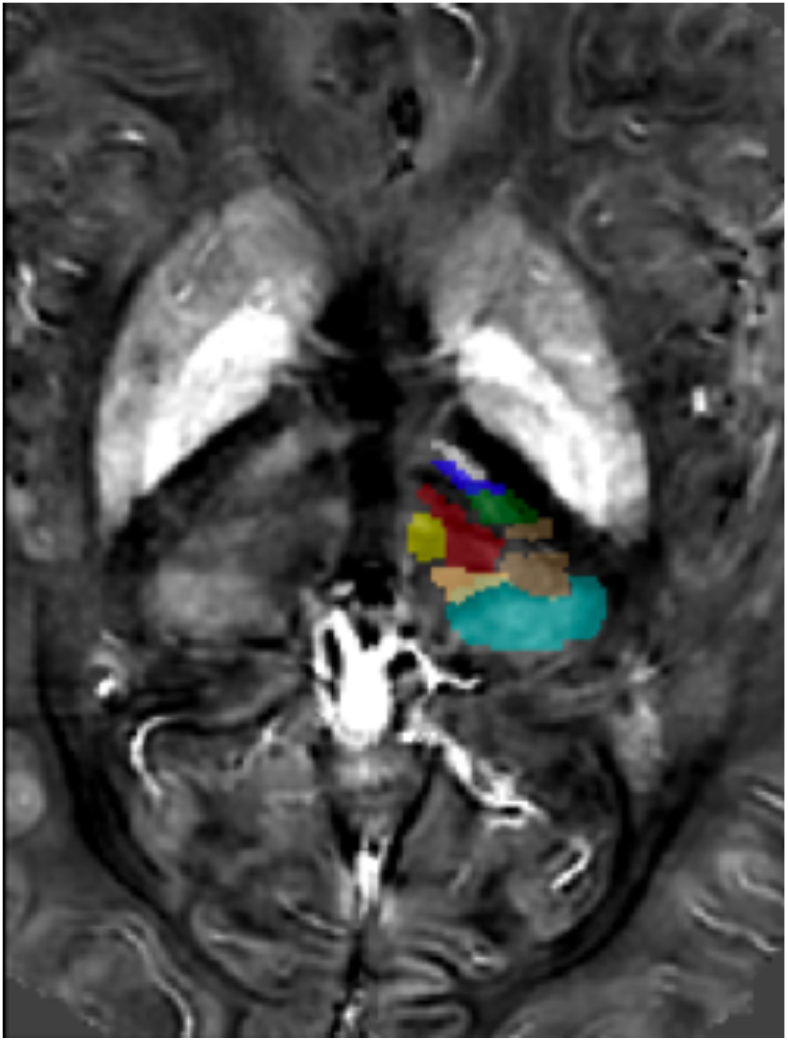
Axial susceptibility map at the level of the mid-thalamus shows multiple substructures within the thalamus, including the pulvinar (cyan), centromedian nucleus (red), ventral lateral anterior nucleus (dark blue), ventral posterior lateral nucleus (beige), ventral intermediate nucleus (green), ventral anterior nucleus (VA - grey), nucleus parafasiculairs (yellow), and nucleus limitans (copper).

Although a variety of acquisition parameters and MRI scanner platforms were utilized, the VIM is consistently visualized as an obliquely-oriented hypointense substructure situated within the mid thalamus, adjacent to the medial margin of internal capsule (Figure 2). Comparing QSM-based direct visualization to the indirect method, similar discrepancies were identified for both the single-orientation and multi-orientation QSM methods. Specifically, the VIM localized indirectly using the Guiot parallelogram was inferior and medial to the location determined using direct visualization (Table 1) (Figure 3). In four of the cases, there was no apparent overlap with the VIM or DRTT using landmark based, indirect method. For two of these cases, localization by the indirect method resulted in a discrepancy of up to 8 mm with respect to the direct targeting approach.

**Table 1:** Average discrepancy (in millimeters) between direct localization of the VIM using QSM and indirect localization using the Guiot parallelogram.

|  | Single-orientation QSM<br>versus Indirect localization<br>(mm) | Multi-orientation QSM<br>versus Indirect<br>localization (mm) | P-value |
| --- | --- | --- | --- |
| Left VIM | $2.7 \pm 0.3$ | $3.4 \pm 0.1$ | $> 0.05$ |
| Right VIM | $4.8 \pm 3.1$ | $4.3 \pm 1.1$ | $> 0.05$ |

**Figure 2:**
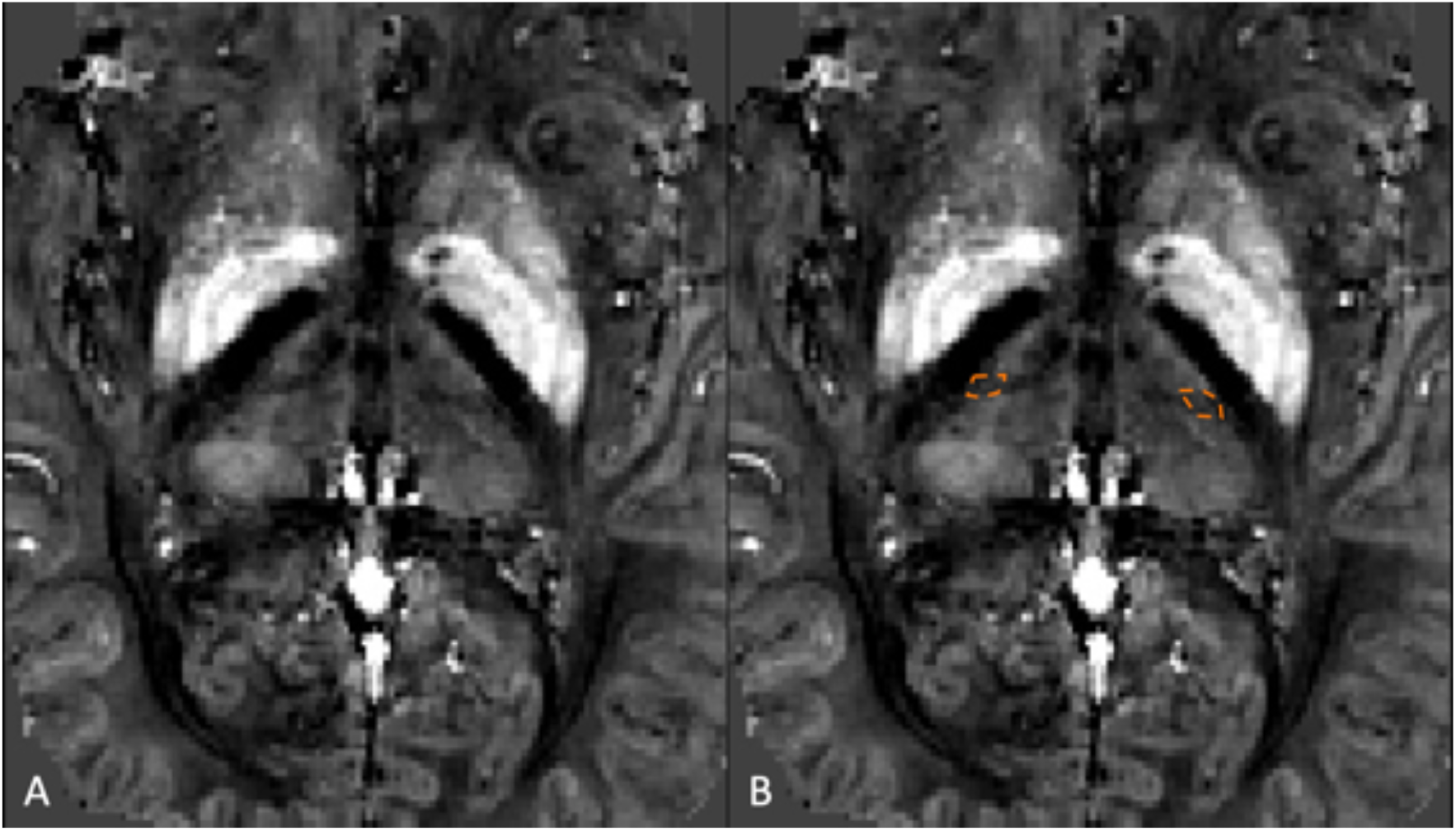
Axial susceptibility map at the level of the thalamus shows the VIM without (A) and with (B) annotation, which appears as a hypointense linear structure within the lateral thalamus situated medial to the posterior limb of the internal capsule.

**Figure 3:**
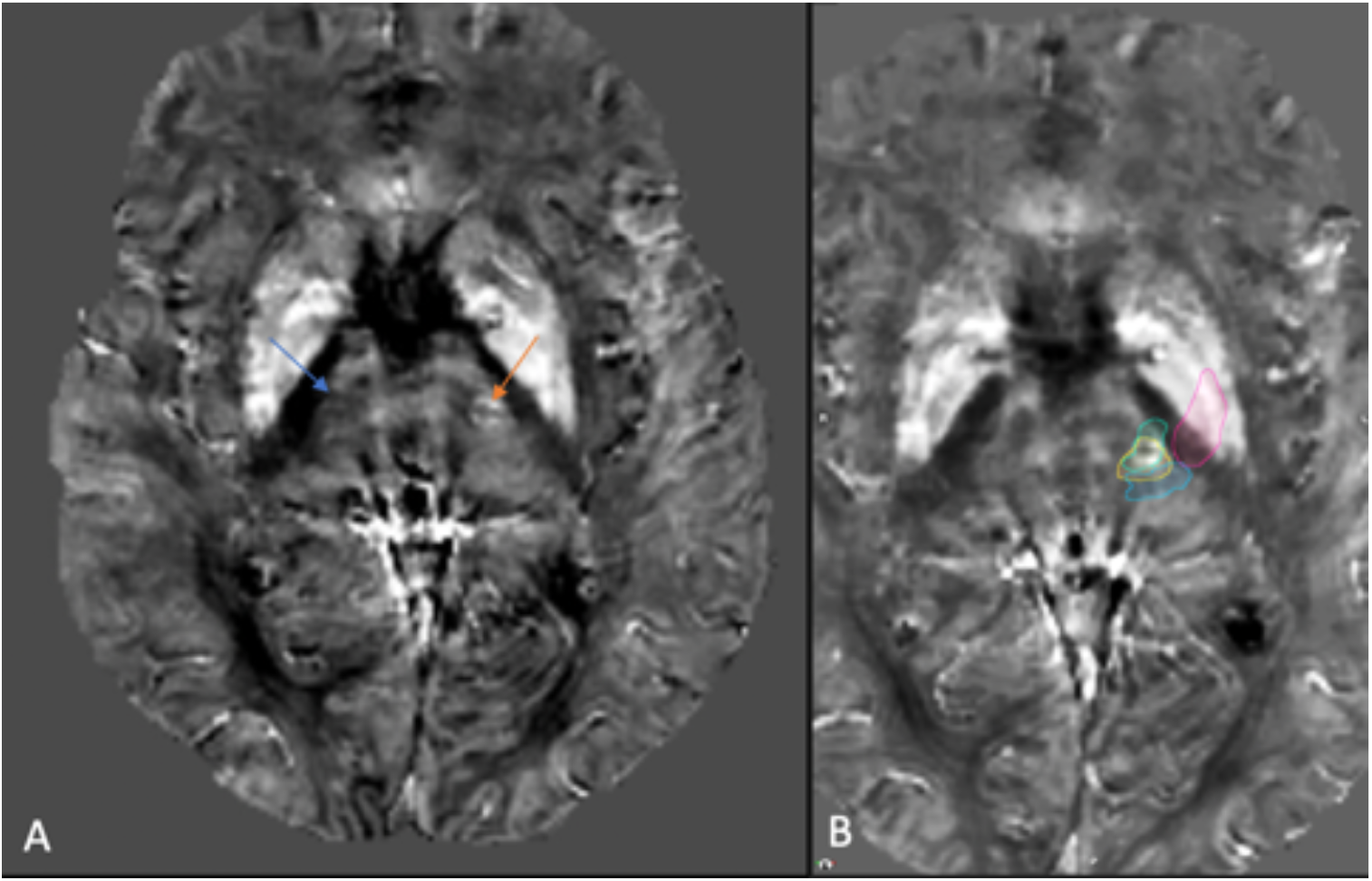
Axial (A) susceptibility map at the level of the thalamus in a patient treated with MRgHIFU within the left VIM (red arrow). The untreated right VIM can be seen (blue arrow). Overlaid diffusion tractography (B) regions showing agreement with the direct localization method (decussating DRTT [green], non-decussating DRTT (yellow], medial lemniscus [blue], and corticospinal tract [red]).

A patient with essential tremor who underwent treatment with MRgHIFU guided by tractography was also imaged. At the site of ablation in the left thalamus, there is focally increased susceptibility signal which corresponds to the location of the VIM based on direct localization (Figure 4a). The post-HIFU QSM demonstrated that the region identified with direct localization coincides with the location of the DRTT (Figure 4b).

**Figure 4:**
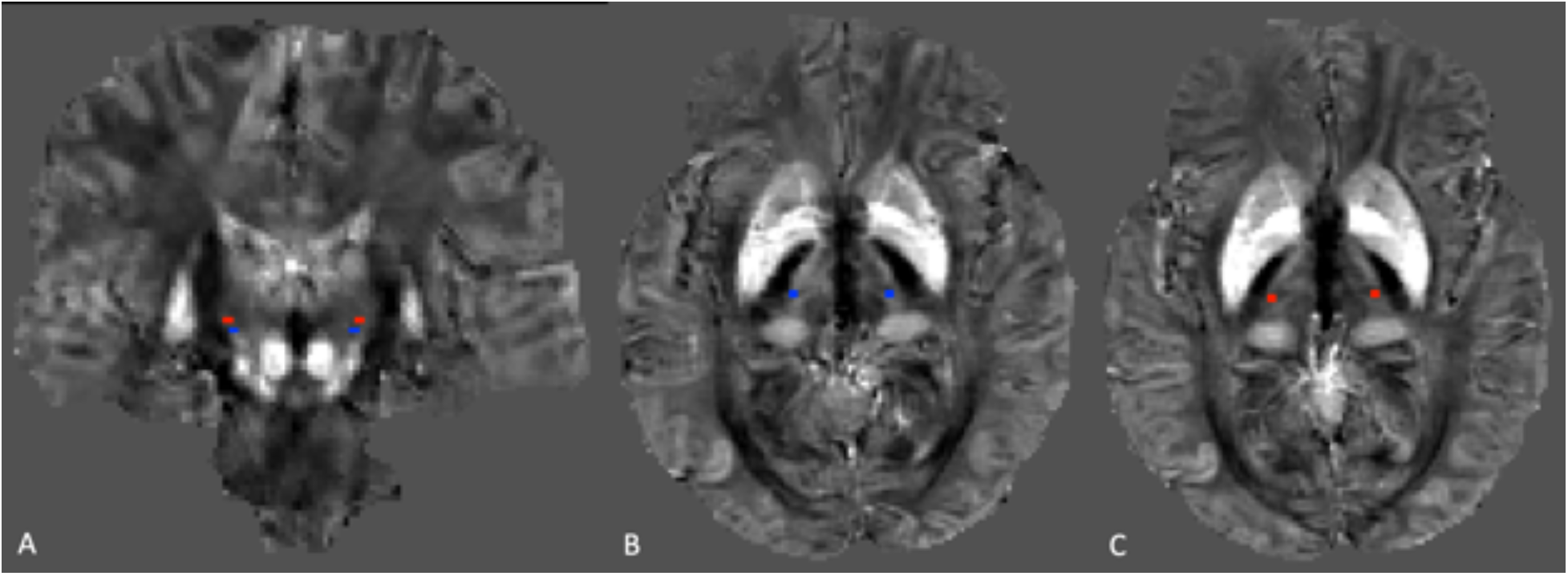
Coronal (A) and axial (B, C) susceptibility maps of the brain demonstrating the discrepancy in localization of the VIM based on the indirect method (blue) and direct method (orange).

## Discussion

In this paper we show that clinically feasible, single-orientation QSM can segment the VIM and other thalamic nuclei based on differences in paramagnetic (iron) and diamagnetic (myelin) content. QSM-based localization of the VIM differs from indirect targeting but correlates well with other advanced imaging techniques such as diffusion tractography.

Direct visualization of the VIM is not possible with standard, high-resolution anatomic MRI sequences. As a result, alternative approaches have been adopted including indirect targeting methods and microelectrode recordings. More recently, there has been renewed interest in developing direct, non-invasive imaging approaches to target thalamic nuclei.(4) For example, the DTI-based approach identifies the DRTT. However, user input is needed for selection of tractography tracking parameters as well as region-of-interest selection, all of which can influence the resultant generated tracts.

To date, there has been only one published study employing QSM to delineate the thalamic nuclei (8). However, that study had important limitations including: 1) a requirement for multiple head positions which is clinically prohibitive 2) the use of 7T MRI, which is not widely available. While these approaches offer certain intrinsic advantages in terms of SNR and circumventing the challenges posed by susceptibility anisotropy, our approach for generating magnetic susceptibility maps from a single orientation scan at 3T appear to offer sufficient contrast to delineate the VIM. It is worth noting that while image quality was higher for the high-resolution single orientation QSM, the VIM could still be readily identified using the 1.1 mm isotropic QSM that could be acquired in approximately half the scan time (6 minutes versus 12 minutes). The disagreement between the anatomic atlas and the expected location of the VIM likely represents a well-described phenomenon in which individual variation is not accounted for by atlas based or indirect targeting methods. This is supported by the fact that in addition to this work other groups have found disagreements between indirect visualization methods and tractography.(3, 16)

Single orientation QSM overcomes several limitations of traditional susceptibility weighed imaging and obviates the need for multiple head positions used in COSMOS QSM. Using single head position multi-echo GRE sequences at 3T to generate magnetic susceptibility maps, we were able to directly visualize the VIM within the lateral aspect of the DRTT, which appeared as a hypointense structure within the lateral aspect of the mid thalamus on the magnetic susceptibility images. The hypointense signal most likely arises from diamagnetic myelin sheaths encasing axons within the DRTT. Furthermore, owing to its high sensitivity to both paramagnetic and diamagnetic tissues including iron and myelin, susceptibility mapping can delineate subtle variations between subcortical substructures that are otherwise invisible on other sequences (8). This allows for visualization of individual thalamic nuclei which can help to guide treatment-planning.

A current limitation of widely adapting QSM is variation in post-processing algorithms which are routinely performed offline instead of at the scanner. However, the consistency with which we were able to visualize the VIM across different of MR scanner settings points to generalizability of the single-orientation QSM approach. Our study also lacked ground truth to further validate our technique. This can be addressed in future cadaver-based studies with histopathologic ground truth or with invasive microelectrode testing. Despite these limitations, single orientation QSM offers a promising advanced imaging technique for visualizing subcortical structures of the brain.

## Conclusion

Single orientation QSM at 3T is able to delineate thalamic nuclei and the DRTT, facilitating localization of the VIM. This approach obviates the need for multiple head positions and ultrahigh field MRI scanners, expanding the clinical potential of QSM in patients with essential tremor.

